# Things softly attained are long retained: Dissecting the Impacts of Selection Regimes on Polymorphism Maintenance in Experimental Spatially Heterogeneous Environments

**DOI:** 10.1101/100743

**Authors:** Romain Gallet, Rémy Froissart, Virginie Ravigné

**Affiliations:** INRA, UMR 385 BGPI, Cirad TA A-54/K Campus International de Baillarguet 34398 Montpellier Cedex 5, France; CNRS, IRD, Université de Montpellier, UMR 5290 MIVEGEC, F-34090 Montpellier, France; CIRAD, UMR PVBMT, F-97410 Saint-Pierre, France

## Abstract

This preprint has been reviewed and recommended by Peer Community In Evolutionary Biology (http://dx.doi.org/10.24072/pci.evolbiol.100020). Predicting and managing contemporary adaption requires a proper understanding of the determinants of genetic variation. Spatial heterogeneity of the environment may stably maintain polymorphism when habitat contribution to the next generation can be considered independent of the degree of adaptation of local populations within habitats (*i.e.*, under soft selection). In contrast, when habitats contribute proportionally to the mean fitness of the populations they host (hard selection), polymorphism is not expected to be maintained by selection. Although mathematically established decades ago, this prediction had never been properly explored. Here we provide an experimental test in which polymorphic populations of *Escherichia* coli growing in heterogeneous habitats were exposed to hard and soft selection regimes. As predicted by theory, polymorphism was longer preserved under soft selection. Complementary tests established that soft selection slowed down fixation processes and could even protect polymorphism on the long term by providing a systematic advantage to rare genotypes.

## Introduction

Genetic variation is the fuel of evolution. Understanding the ultimate forces that shape the amount of genetic variation within populations is therefore a central issue of evolutionary biology. Beyond its fundamental interest, this topic is also crucial for a number of applied issues where evolutionary potential matters. In conservation biology for instance, preserving the adaptive potential of endangered species is now a primary goal of management policies (Crandall *et al.* 2000). Similarly, as pathogen evolution regularly ruins management attempts (*e.g.,* antibiotic resistance, plant resistance breakdown), managing pathogen polymorphism is becoming a growing concern (Vale 2013).

The spatial heterogeneity in selection pressures among the different habitats composing an environment constitutes a good explanation of the huge amount of genetic variation observed *in natura*. Yet theoretical works have previously shown that spatial heterogeneity does not necessarily lead to the stable maintenance of local adaptation polymorphism (Dempster 1955, Christiansen 1974, de Meeûs *et al.* 1993, see Kassen 2002, Ravigné *et al.* 2009, Massol 2013, Vale 2013 for reviews). Whether selection leads to the stable maintenance of diversity depends on the interaction between several factors; the existence and strength of local adaptation trade-offs (*i.e.* negative genetic correlations in fitness across different habitats, Levins 1962), the frequency and productivity of the different habitats in the environment (Levene 1953), and the amount of gene flow between habitats (Maynard Smith 1966, Christiansen 1975, Débarre and Gandon 2011). Moreover, very early models showed that the possibility for stable polymorphism crucially depended on how local populations within the different habitats contribute to the next generation (Levene 1953, Dempster 1955, Christiansen 1975, Maynard Smith and Hoekstra 1980). In some organisms, the contribution of local populations to the next generation is fairly independent of their genetic composition. Density regulation limits the productivity of local populations so that better adaptation to their habitat does not translate into higher local productivity. This can be observed in solitary insect parasitoids that can only lay one egg per host individual (Mackauer 1990). In pathogens, within-host pathogen accumulation is thought to increase the risk of premature host death. This mechanism limits transmission between hosts. In this case, pathogens infecting the same host are a local population, so that the density regulation of the contribution of the population is suspected to be the rule rather than the exception (de Meeûs *et al.* 1998, Chao *et al.* 2000). This type of density regulation produces a selection regime called soft selection (Levene 1953, Wallace 1975) that has theoretically been shown to be prone to diversification and polymorphism maintenance.

In contrast, in other species and environments under hard selection, habitat contribution to the next generation is not a fixed characteristic but rather depends on the genetic composition of the local population, *i.e.*, better adaptation implies greater habitat contribution to the next generation (Dempster 1955, Wallace 1975). Hard selection, in principle, hampers diversification and polymorphism maintenance and is expected when population density is not regulated locally within each population but globally at the scale of the environment. It can also be observed in cases where adaptation increases the carrying capacity of the habitat through, *e.g.,* a more efficient use of nutrients. In the case of pathogens for instance, this type of selection occurs when transmission does not depend on whether the host is dead or alive. Hence hard selection is likely frequent in serial passage experiments when parasite transmission is simulated by experimenters. Logically, most serial passage experiments lead to a decrease or disappearance of the initially present polymorphism (for review, see Ebert 1998).

Despite a vast consensus among theoreticians over the importance of the selection regime for polymorphism maintenance in heterogeneous environments, the concepts of hard and soft selection generally remain overlooked in the empirical literature. Hard and soft selections have recently become an explicit concern in studies measuring selection strength and mutation accumulation (Juenger *et al.* 2000, Kelley *et al.* 2005, Laffafian *et al.* 2010, Wade *et al.* 2010, Johnson *et al.* 2011). Despite their relevance, the terms hard and soft selection are still not mentioned in many fields where it could be important for analyses to distinguish between these selection regimes. For example, these concepts could be particularly useful for understanding plant pest evolution in landscapes composed of mixtures of plant varieties or the evolution and management of antibiotic resistance.

It must be recognized that despite several important attempts (*e.g.,* Bell 1997), proof of concept – through a proper experimental test – has yet to be made (Vale 2013). Some experiments did test the effect of spatial heterogeneity on genetic variability (reviewed in Rainey *et al.* 2000, Kassen 2002, see also Jasmin and Kassen 2007), most of them concluding that populations confronted with a spatially heterogeneous environment are more variable than those exposed to homogeneous environments. Yet, these experiments did not control for the selection regime imposed by serial passages and experimentally applied hard selection (but Garcìa-Dorado *et al.* 1991, Bell and Reboud 1997). The higher variability observed under these heterogeneous treatments admittedly lied in transient polymorphism being less efficiently removed from heterogeneous environments than from homogeneous environments. A possible exception was the experiment by Bell and Reboud (1997) in which, despite no experimentally-imposed density regulation, local density regulation was suspected to have occurred and to have promoted higher genetic variance in heterogeneous environments as compared to homogeneous ones. One study explicitly imposed hard and soft selection regimes on a mixture of strains of the unicellular algae *Chlamydomonas reinhardtii* maintained in a heterogeneous environment for 50 generations without sexual reproduction (Bell 1997). Contrary to theoretical predictions, genetic variation remained similar regardless of the type of density regulation. This unexpected result was interpreted as a consequence of the specific nature of the environmental heterogeneity – habitats were composed of different mixtures of nutrients – that did not impose a trade-off in local adaptation (Bell 1997). In the absence of such trade-offs, even under soft selection, polymorphism is not selected for.

Here we aimed at experimentally testing the prediction that soft selection can produce the negative frequency-dependence required for stable maintenance of polymorphism (Karlin and Campbell 1981). To create the local adaptation trade-off required for polymorphism maintenance, polymorphic *Escherichia coli* populations were built using two genotypes, one being resistant to tetracycline and the other to nalidixic acid. These populations were grown in three heterogeneous environments each composed of two different habitats, one containing a very low concentration of tetracycline and the other a very low concentration of nalidixic acid. Low antibiotic concentrations provided a selective advantage to the resistant genotype over the susceptible one, but both genotypes could grow in all conditions. Three different trade-offs were produced by varying habitat productivities. As in Bell (1997), serial passages were controlled to apply either hard selection (*i.e.*, by transferring an aliquot of each environment) or soft selection (*i.e.*, by transferring a fixed number of cells from each environment). The evolution of genotype frequencies over the course of the experiment was precisely monitored and systematically compared to theoretical predictions obtained assuming either hard or soft selection. The duration of the experiments was kept short enough to avoid the emergence (by *de novo* mutation) of a generalist genotype. Whether observed polymorphisms could be maintained on the long term was checked *a posteriori*, through a complementary experiment testing for polymorphism protection. A polymorphism is said ‘protected’ if the genotypes involved increase in frequency when rare. Under such conditions, none of the genotypes can ever be eliminated from the population and the polymorphism can be maintained as long as the trade-off exists. In all conditions tested polymorphism was either lost during the experiment or bound to disappear (it was not protected) under hard selection. In contrast in the same conditions, a pulse of soft selection once every few generations was sufficient for polymorphisms to be maintained during the experiment and to be protected by negative frequency-dependent selection. We discuss the potential of such experimental system to explore the contribution of soft and hard selection to local adaptation polymorphisms.

## Materials and Methods

### Bacterial strains

The *E. coli* B strains used in this study, REL4548 YFP-Tet^R^ and REL4548 CFP-Nal^R^ derive from the strain REL4548 kindly provided by R. E. Lenski. REL4548 was evolved in Davis minimal (DM) medium supplemented with 25 μg/mL glucose (DM25) for 10,000 generations as part of a long-term evolution experiment (Lenski *et al.* 1991). Gallet *et al.* (2012) then inserted YFP and CFP genes at the *rhaA* locus of REL4548 using a technique developed by Datsenko and Wanner (2000). A mini-Tn*10* derivative 104 — which contains a tetracycline resistance cassette (Kleckner *et al.* 1991) — was introduced at the *insL-1* locus into REL4548 YFP (clone T121) (Gallet *et al.* 2012) to construct REL4548 YFP-Tet^R^. The strain REL4548 CFP-Nal^R^ was then created by selecting a resistant REL4548 CFP colony on a LB plate (10 g/L NaCl, 10 g/L tryptone, 5 g/L yeast extract; 15 g agar, 1000 mL H2O) supplemented with 20 μg/mL of nalidixic Acid. These constructions permitted the association of a specific antibiotic resistance with a specific fluorescent marker and therefore easily identifying resistant strains by their fluorescence. Bacterial strains were stored at -80°C in 15 % glycerol stocks.

### Habitats

Four habitats (*i.e.*, growth media) were used. Each habitat hosted a single local population. They differed in productivity (*i.e.,* glucose concentration in growth medium), and/or by the presence of very low concentrations of either tetracycline or nalidixic acid. All media were made on the base of Davis minimal (DM) medium (KH2PO4 monohydrate 5.34 g/L, KH2PO4 2 g/L, ammonium sulfate 1 g/L, sodium citrate 0.5 g/L). Bottles were weighted before and after autoclaving and sterile milliQ water was added to compensate for evaporation happening during sterilization. After autoclaving, media were supplemented with 806 μL/L of MgSO_4_^2-^ [1 M], 1 mL/L Thiamine (vitamin B1) [0.2%]. Then, 40 μL/L or 1 mL/L of glucose [2.5%], were added in order to make DM2 and DM50 (2 and 50 μg/mL of glucose being present in the medium, respectively). These media were equivalent to the one used by Lenski *et al.* (1991), but with different glucose concentrations. Antibiotics were used at subinhibitory concentrations to provide a moderate fitness advantage to the resistant genotype. To take into account week-to-week variations (different medium batch, antibiotic dilution etc.), culture media were tested prior to start the experiments, and the relative fitnesses of bacterial genotypes were measured. Antibiotic concentrations were adjusted in order to reach similar fitness in replicated experiments. Thus, tetracycline and nalidixic acid were added at final concentrations of 0.02 μg/mL and 0.7 μg/mL respectively for the first experiment and 0.03 μg/mL and 0.8 μg/mL respectively for the second experiment (which resulted in similar relative fitness in both experiments). Habitats were hereafter denoted Nal2, Nal50, Tet2, and Tet50 depending on the antibiotic used and their productivity as measured through DM concentration.

### Environments

Three different environments were used, each composed of two habitats (Figure 1A). These three different environments correspond to three different local adaptation trade-offs. In one environment, habitat productivities were comparable (environment B in figure 1B, composed of Nal50 and Tet50 habitats, hereafter referred to as ‘symmetric’ environment). In the two other environments (hereafter ‘asymmetric’ environments), one habitat was more productive than the other (environment A on figure 1B was composed of Nal2 and Tet50 habitats and environment C was composed of Nal50 and Tet2 habitats). While in environment B, both bacterial genotypes had similar mean fitness in the habitat they were adapted to, environments A and C led to favor one genotype over the other at the scale of the whole environment (Figure 1B).

**Figure 1:**
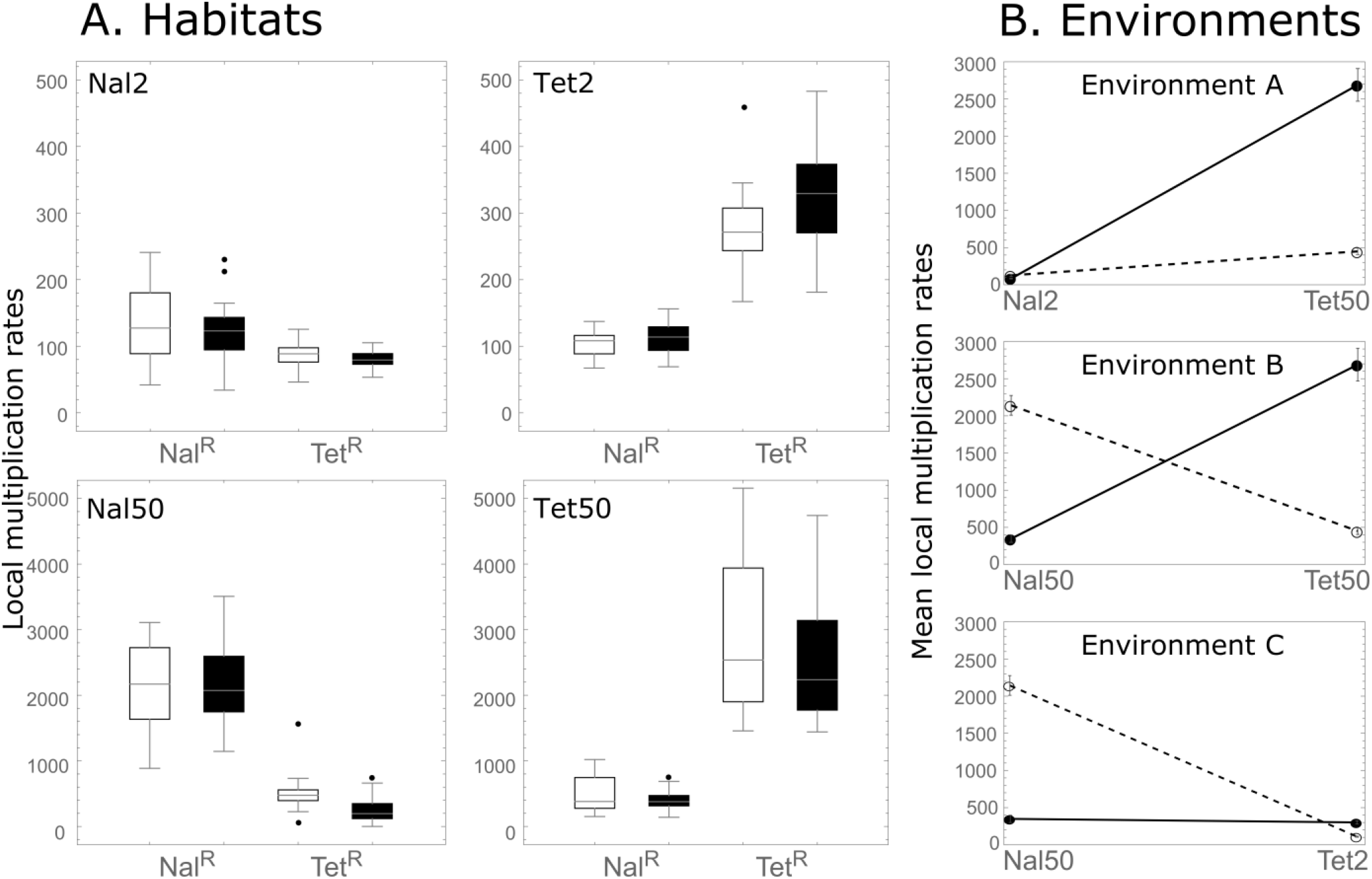
Habitats, environments and local adaptation trade-offs. Local fitness is defined as the rate of multiplication within one habitat between transfers (thereby corresponding to viabilities in Levene’s model). Panels: A. Distributions of the local multiplication rates of Nal^R^ and Tet^R^ genotypes in the long (white whisker charts) and the short (black whisker charts) trials of Experiment 1 for the four habitats used. B. Mean local multiplication rates of Nal^R^ (open circles and dashed line) and Tet^R^ (filled circles and plain line) genotypes in the three environments used (with confidence intervals over all replicates and transfers of Experiment 1).

### Experiment 1: Maintenance of established polymorphism

In a first experiment (hereafter referred to as Experiment 1, figure 2), polymorphic populations with initially equal frequencies of both genotypes were grown under hard selection and soft selection regimes. For each of these two selection regimes, three replicate populations were used for each of the three environments (3 replicates × 3 environments × 2 selection regimes = 18 populations in total). Before the start of the experiment, REL4548 YFP-Tet^R^ and REL4548 CFP-Nal^R^ genotypes were grown separately overnight in 5 mL of DM25 (37°C, 215 rpm). At T0 the optical density (OD, 600nm, Eppendorf spectrophotometer) of each culture was measured and a 50/50 mix was made to inoculate all habitats of all environments. At the end of each day (Figure 2), a starting bacterial population was prepared by mixing the bacterial populations from the two habitats. Depending on the selection regime, either the same volume (50 μL – hard selection) or different volumes (containing 10^7^ cells per habitat – soft selection) where added to the mix, with a 10-fold dilution in DM0 (*i.e.,* DM medium containing no glucose). Part of the mix was used to make a glycerol stock (stored at -80°C) for subsequent flow cytometer analysis while the other part was used to inoculate both habitats of the environment of the next passage (50 μL into 5 mL of fresh media – an additional 100fold dilution). Populations were grown overnight (37°C, 215 rpm, 18h of incubation). The whole experiment was replicated twice. In the first trial, flow cytometer measurements showed that the realized initial frequency of REL4548 YFP-Tet^R^ was 0.508 and the experiment was conducted over five transfers. The second trial, conducted simultaneously with Experiment 2, started from an initial frequency of REL4548 YFP-Tet^R^ of 0.437 and had to be interrupted after three transfers due to technical difficulties.

**Figure 2.**
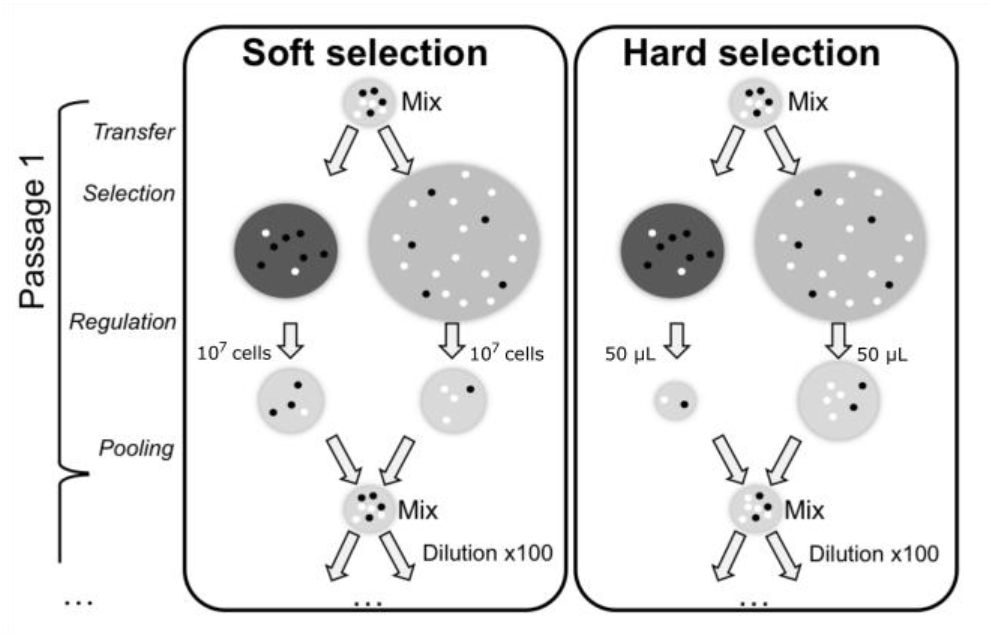
Experimental design. Light and dark grey circles represent habitats supplemented with tetracycline and nalidixic acid respectively. Circle sizes illustrate the nutrient richness of these habitats (large circles = DM50, small circles = DM2). This figure illustrates environment C (Tet2-Nal50). “Mix” grey circles represent mixing tubes. Tiny black and white circles represent Tet^R^ and Nal^R^ cells respectively. Black cells have a selective advantage in the dark grey environment, while white cells have a selective advantage in the light grey environment. Populations were transferred daily for 5 transfers. The experiment starts with the transfer of a 50/50 Nal^R^ - Tet^R^ mix in each habitat. During the selection step of the experiment, cells grow in the habitat. The amount of cells transferred during the regulation step depends on the selection treatment. Under hard selection fixed volumes (50 μL) of each habitat were pooled together in the mixing tube, while under soft selection fixed numbers of cells (10^7^ cells) from each habitat were pooled.

### Experiment 2: Polymorphism protection

To go further and test the hypothesis that soft selection can maintain genetic diversity indefinitely by producing negative frequency-dependence, while hard selection cannot, we conducted a complementary experiment (Experiment 2) testing for polymorphism protection. Polymorphism is protected by negative frequency-dependent selection if and only if both genotypes increase in frequency when rare (Prout 1968). In other words, this experiment allows showing that as long as the trade-off persists the genotype with the lowest absolute fitness will not be eliminated from the population, thereby fostering the maintenance of polymorphism. We therefore applied hard and soft selection on initial populations where the genotype with a global disadvantage in the considered environment was rare. In Environment A (Nal2-Tet50) the initial frequency of REL4548 YFP-Tet^R^ was 0.975. In Environment C (Nal50-Tet2) the initial frequency of REL4548 YFP-Tet^R^ was 0.035. In Environment B (Nal50-Tet50), initially conceived as symmetric, both initial frequencies were tested. The experiment was conducted over two transfers only.

### Flow cytometry

Flow cytometry was performed on a Gallios flow cytometer (Beckman Coulter Inc) designed to detect small objects such as bacteria. We used flow cytometry to estimate (*i*) the relative genotype frequencies and (*ii*) cell concentration. This procedure was performed on overnight cultures and on mixes. To estimate cell concentration, fluorescent beads of known concentrations (AccuCount Fluorescent Particles, 7.0-7.9 μm, Spherotech) were added to the cells. Results were analyzed with the Kaluza 1.3 software (Beckman coulter Inc).

### Local fitness measurements

The local fitnesses of the two bacterial genotypes in the four habitats were measured in two complementary manners. Firstly, using flow cytometry, for each habitat, the rate of multiplication of each genotype between two transfers was computed as the ratio of cell concentration at the end of the overnight culture over cell concentration at the beginning. These multiplication rates are akin to viability coefficients as defined in hard selection models and could therefore be directly used to feed Levene’s and Dempster’s equations to establish theoretical predictions. Secondly, for the sake of comparison with other works on bacteria, selection coefficients (*sensu* Chevin 2011) were calculated:

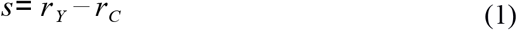

with

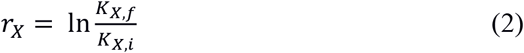

where *K*_*X,i*_ and *K*_*X,f*_ are respectively the initial and final effective of genotype X. Selection coefficients, available in Appendix S1, were used to confirm that no evolution towards a generalist phenotype was observed during the experiment.

### Theoretical predictions

Given the viability coefficients *W*_*i,j*_ of genotype *i* in habitat *j*, under soft selection, the change in frequency *p*_*t*_ of Tet^R^ bacteria from transfer *t* to transfer *t* + 1 is governed by the following equation:

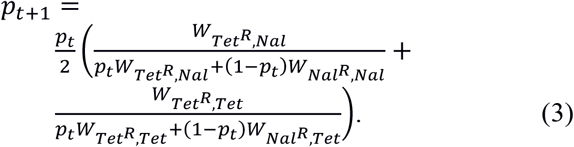

Under hard selection, the trajectory of the frequency *p*_*t*_ of Tet^R^ bacteria is given by:

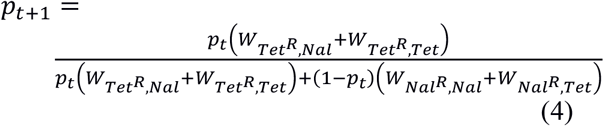

In principle it is thus possible to compare the observed trajectories of genotype frequencies to theoretically expected ones under both selection regimes. Local fitnesses imposed by habitats to the two genotypes were experimentally variable (Figure 1A). To account for such experimental variability, 10,000 trajectories of Tet^R^ frequency over transfers were simulated by random sampling, at each transfer, of the values of viability coefficients 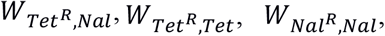, and 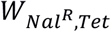, among all corresponding values observed over all transfers of Experiment 1 for each habitat. The median and 2.5^th^ and 97.5^th^ percentile values of the distribution of Tet^R^ frequency at each transfer were used to represent the theoretically expected trajectories of genotype frequency. Similarly, the equilibrium Tet^R^ frequency was estimated using the median, 2.5^th^ and 97.5^th^ percentile values of the distribution of predicted Tet^R^ frequencies after 100 transfers.

## Results

### Three heterogeneous environments with clear local adaptation trade-off

Figure 1A shows the local fitnesses (*i.e.,* between transfer multiplication rates) obtained for each bacterial genotype over all replicates in each of the four habitats. From this and the computation of selective coefficients (available in appendix S1), the existence of three different local adaptation trade-offs could be verified (Figure 1B). It was also confirmed that local fitnesses were similar in the two independent trials of Experiment 1 and that multiplication rates – hence bacteria – did not evolve during the experiment (Figure S1). Theoretical predictions showed that Environment A (Nal2-Tet50) was so asymmetric that the fixation of the Tet^R^ genotype was expected under both hard and soft selection (Figure 3A and B, right hand side of the x-axis). The expected dynamics of genotype frequency however differed clearly between hard and soft selection (grey areas in figure 3A and B). In Environments B (Nal50-Tet50) and C (Nal50-Tet2), soft selection was expected to lead to polymorphism maintenance (Figure 3D and F), while hard selection was expected to lead to the fixation of one of the two genotypes (Tet^R^ in Environment B and Nal^R^ in Environment C, figure 3C and E). In Environment B, the dynamics of genotype frequencies over 5 transfers were hardly distinguishable between hard and soft selection (Figure 3C and D).

**Figure 3:**
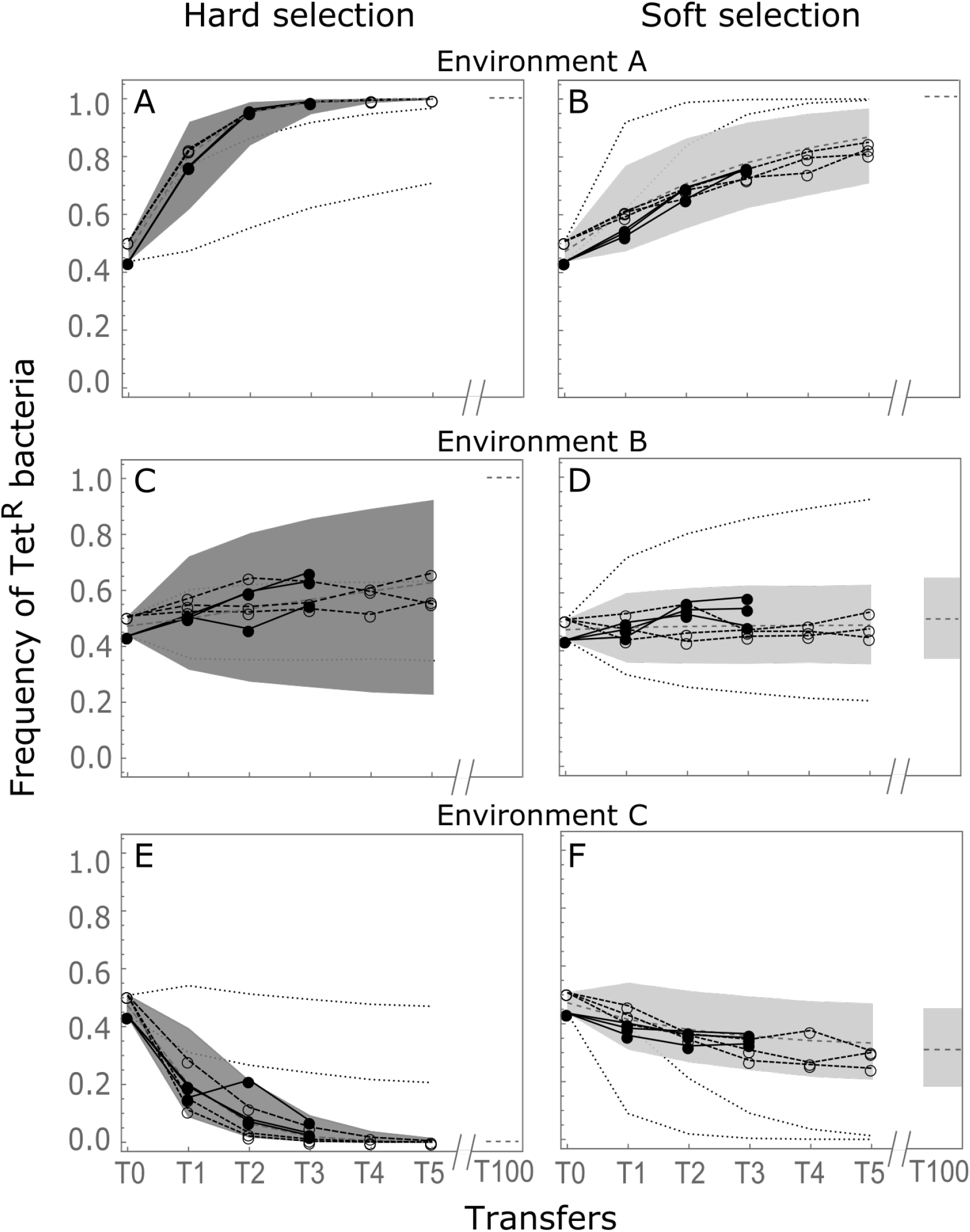
Evolution of genotype frequencies under hard and soft selection. All panels show the dynamics of the frequency of Tet^R^ bacteria over successive transfers in Experiment 1. Filled circles and lines show the frequencies observed in the three transfers of the short trial. Open circles and dashed lines correspond to the five transfers of the long trial. Note that initial frequencies slightly differ between the two trials. Grey areas show the 95% envelopes of theoretically predicted frequencies under the corresponding selection regime. Dashed grey lines show the medians of theoretical frequencies. Dotted black lines delimit the 95% envelopes of the other selection regime (e.g. on the hard selection panels, dotted lines show the predictions under soft selection). At the right of the x-axis break, theoretically predicted equilibrium frequencies are shown. Panels: A. Environment A (Nal2-Tet50) under hard selection. B. Environment A under soft selection. C. Environment B (Nal50-Tet50) under hard selection. D. Environment B under soft selection. E. Environment C (Nal50-Tet2) under hard selection. F. Environment C under soft selection.

### Effects of selection regimes on the maintenance of polymorphism

Under hard selection, in both environments with asymmetric habitat productivities (Environments A - Nal2-Tet50 and Environment C - Nal50-Tet2), polymorphism was almost completely lost over the experiment (Figures 3A and E). In environment A, Tet^R^ genotype frequency reached an average of 0.991 ± 0.001 after 3 transfers in the two Experiment 1 trials (n=6 replicates) and 0.999 ± 0.0002 after 5 transfers in the long trial (n=3). In environment C, Tet^R^ genotype frequency decreased to 0.032 ± 0.027 after 3 transfers in the two Experiment 1 trials (n=6) and 0.002 ± 0.003 after 5 transfers in the long trial (n=3). The trajectories of genotype frequencies fit well with predictions obtained assuming hard selection (dark envelop of theoretical predictions obtained for soft grey in figures 3A and E) and fell outside the 95% envelop of theoretical predictions obtained for soft selection (dotted lines in figures 3A and E). In the symmetric environment (Environment B), as predicted, polymorphism was almost unchanged at the end of the experiment with only a slight increase of TetR frequency. On average, TetR genotype frequency reached 0.595 ± 0.055 after 3 transfers in the two trials of Experiment 1 (n=6 replicates) and 0.593 ± 0.059 after 5 transfers in the long trial (n=3).

Under soft selection, genetic polymorphism was maintained throughout the experiment regardless of habitat productivities (Figures 3B, 3D and 3F). In Environment A (Nal2-Tet50), the frequency of TetR bacteria increased at a rate compatible with predictions obtained under soft selection (light grey area in figure 3B) and not with predictions obtained under hard selection (dotted lines in Figure 3B). In this environment, although the expected final outcome of selection was the same under hard and soft selection regimes (fixation of TetR bacteria), the rate of evolution was much slower under soft selection than under hard selection.

In Environments B and C where stable polymorphism was expected, genotype frequencies fit well with predictions obtained under soft selection (light grey areas in figures 3D and F). In the five-transfer trial, the frequency of Tet^R^ bacteria finally attained 0.484 ± 0.045 (expected value: 0.488 with 95% envelope [0.353-0.628]) in Environment B and 0.284 ± 0.033 (expected value: 0.333 with 95% envelope [0.206-0.470]) in Environment C.

### Polymorphism protection observed in soft but not in hard selection

The trajectories of genotype frequency observed in Experiment 2 were again in agreement with theoretical expectations (Figure 4). In Environment A (Figure 4A), as observed previously, neither hard selection nor soft selection produced an advantage for the rare genotype. In Environment B the Tet^R^ genotype had a global fitness advantage over the Nal^R^ competitor (Figure 4C). Tet^R^ frequency nevertheless significantly decreased from a high initial starting value under soft selection. Similarly in Environment C (Figure 4D), where Tet^R^ bacteria had a global fitness disadvantage, Tet^R^ frequency significantly increased when initially rare under soft selection only. In these two environments the observed polymorphism was therefore protected under soft selection.

**Figure 4:**
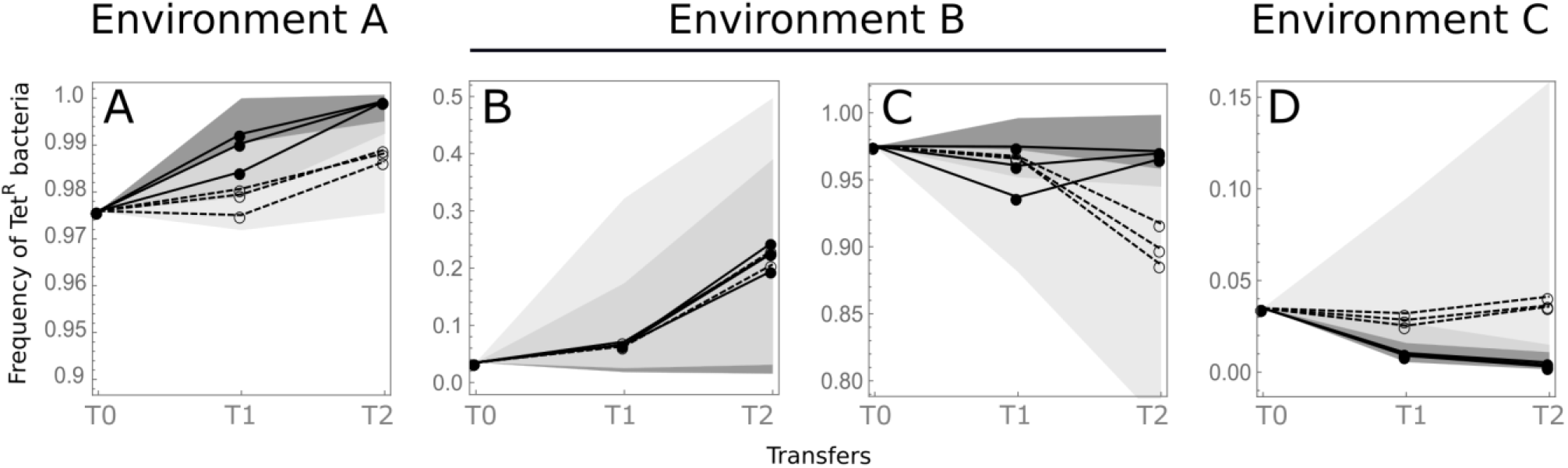
Polymorphism protection. All panels show the dynamics of the frequency of Tet^R^ bacteria over successive transfers in Experiment 2. Filled circles and lines show the frequencies observed under hard selection. Opened circles and dashed lines correspond to soft selection. Dark and light grey areas show the 95% envelopes of theoretically predicted frequencies under hard and soft selection, respectively. Panels: A. Environment A (Nal2-Tet50) with initially frequent Tet^R^ bacteria. B. Environment B (Nal50-Tet50) with initially rare Tet^R^ bacteria. C. Environment B with initially frequent Tet^R^ bacteria. D. Environment C (Nal50-Tet2) with initially rare Tet^R^ bacteria.

## Discussion

From the seminal debate between Levene (1953) and Dempster (1955), theory has suggested that the way populations redistribute among habitats of a given environment is crucial for the long-term maintenance of local adaptation polymorphisms. Under some conditions, soft selection, in which habitat contribution to the next generation is constant, can protect polymorphism by producing negative frequency-dependent selection. In contrast, in the same conditions, hard selection – in which habitat contribution to the next generation varies with habitat genetic composition – does not so (*e.g.,* Christiansen 1974, 1975, Karlin and Campbell 1982, de Meeûs *et al.* 1993). Although firmly established this prediction still remains to be experimentally investigated. In the present study hard and soft selection were applied to populations composed of two bacterial genotypes in heterogeneous environments composed of two habitats and whether the initial polymorphism was maintained or not was monitored.

Studying several well-characterized reproducible trade-offs was an important condition of the study. In the absence of a local adaptation trade-off, selection is expected to favor a single generalist genotype and environmental heterogeneity cannot lead to stable polymorphism maintenance. The absence of a clear local adaptation trade-off had led previous experimental tests of hard and soft selection to inconclusive results (reviewed in Vale, 2013). Here, bacterial genotypes and habitats were designed using antibiotic resistance so that each genotype was locally adapted to one habitat. The use of very low antibiotic concentrations was crucial. High antibiotic concentrations would have completely inhibited the growth of susceptible competitors, while very low concentrations simply provide a small fitness advantage to the resistant genotype. Although both genotypes could develop in both habitats, each genotype was specialist of one habitat. The considered trade-off was however not constitutive of bacterial metabolism, and it could have been circumvented by evolution of nalidixic acid resistance in Tet^R^ resistant bacteria for instance. This prevented the use of this experimental design on the long term, the selection of a generalist genotype being avoided by keeping the experiments short enough (≈50 generations). It was further verified that no generalist genotype evolved by monitoring genotypes coefficient of selection throughout the experiment.

This important requirement also implied that long term monitoring of the polymorphism was impossible. This said, in nature many polymorphisms can be observed over long periods of time, even in absence of a clear local adaptation trade-off. Two underlying causes can be identified. First, transient polymorphism is less efficiently removed under soft selection than under hard selection (as observed in Bell, 1997, Jasmin and Kassen 2007). Second, negative frequency-dependence caused by factors other than environmental heterogeneity could be at work (*e.g.,* Hori 1993, Sinervo and Lively 1996, Gigord *et al.* 2001, Olendorf *et al.* 2006). Therefore observing polymorphism maintenance is not sufficient to conclude that environmental heterogeneity selects for polymorphism. Instead, discerning whether observed polymorphisms are due to the negative-frequency dependence produced by environmental heterogeneity or not is of prime importance. To do so, one has to test for polymorphism protection. Polymorphism is protected if both genotypes increase in frequency when initially rare. Under polymorphism protection, no genotype can ever disappear and the polymorphism is bound to maintain (unless important drift causes the random loss of one genotype). Polymorphism protection was evidenced by performing the experimental evolution starting with a population where the genotype with the lowest fitness at the scale of the environment was rare (< 3%) and monitoring genotype frequencies over a few generations (Experiment 2). This second experiment confirmed that the polymorphism already observed during 50 generations in Experiment 1 was expected to be maintained as long as the trade-off exists. As environmental parameters, including habitat quality, were the same in hard and soft selection treatments, it further showed that the mechanism at work was negative frequency-dependent selection imposed by environmental heterogeneity.

In all treatments, experimental results showed a remarkable similarity to theoretical predictions. Under the conditions investigated, hard selection never protected polymorphism. The fixation of the genotype with the highest mean fitness at the scale of the environment was observed within 3 transfers in the two asymmetric environments. In the symmetric environment, polymorphism was still observed after 5 transfers under hard selection. But deviations to frequencies theoretically expected under soft selection (Figure 3) and the polymorphism protection experiment (Figure 4) confirmed that such polymorphism consisted of transient polymorphism not being easily removed because of very similar initial frequencies and local fitnesses. In contrast, under soft selection, polymorphism was never lost over the course of the experiment, even in asymmetric environments where the low fitness genotype was inoculated at a very low relative frequency. This was verified using theoretical predictions and the complementary experiment that such polymorphism was only transient in one of the two asymmetric environments (Environment A), and that it was effectively protected by the existence of a systematic advantage of the rare (*i.e.*, negative frequency-dependence) in the two other environments (Environment B and C). In these situations, even though one genotype has a higher mean fitness at the scale of the environment, the local regulation step that occurs at each transfer opposes the effect of within-habitat selection and hampers invasion of the whole environment by the genotype adapted to the most productive habitat (Levene 1953). Interestingly, in the environment C under soft selection (Figure 3F), the genotypic relative frequencies seem to have reached the equilibrium point predicted by theory. Lastly the experiment confirmed that with all else being equal, even when soft selection is expected to lead to the fixation of a single genotype (*i.e.,* when its mean fitness at the scale of the environment is very high – environment A in figure 3), soft selection leads to a slower rate of evolution than hard selection (as shown by *e.g.,* Whitlock 2002).

The present experiment is a proof of concept seeking for conditions under which the maintenance of a local adaptation polymorphism can be attributed to soft selection. It departed from real-world dynamics by using engineered bacteria in controlled environments. It also departed from the mathematical models of hard and soft selection. In these, population mixing and redistribution among habitats are assumed to happen at each generation and local selection within habitats is assumed to act on viabilities, *i.e.*, either fecundities or survival rates. Here, transfers were controlled to reproduce the density-regulation steps characteristic of hard and soft selection. But between transfers, population growth processes (including birth and death) within environments were left uncontrolled over 8 to 10 generations per transfer. Nothing impeded the occurrence of complex population dynamics or density-dependence within habitats. Bacterial populations could for instance reach their carrying capacity before transfers, so that density regulation could be at work within habitats. The present experiment therefore confirmed that the effects of hard and soft selection at the whole-environment scale were robust to local dynamics. It also suggests that a density regulation step (*i.e.*, transfers in the present experiment) once every few generations is sufficient to trigger polymorphism protection.

From a theoretical perspective it is understood that the conditions for polymorphism maintenance under soft selection are rather stringent (Prout 1968, Christiansen 1974, Maynard Smith and Hoekstra 1980). Various processes, such as drift and mutation, may reduce the range of parameters (trade-off shapes and habitat frequencies) where polymorphism is protected. This suggests that soft selection may not be that frequent in nature and that most observed polymorphism is either transient or maintained by other frequency-dependence mechanisms (de Meeûs *et al.* 2000). To some extent, the present study contradicts this view and suggests that the importance of soft selection in shaping standing genetic variation should not be overlooked (Agrawal 2010, Reznick 2016). In recent experiments, the ‘softness’ of selection (*i.e.,* the contribution of soft selection) was measured in experimental populations of both *Drosophila melanogaster* with different genes (Laffafian *et al.* 2010, Ho and Agrawal, 2014) and on seedling emergence in *Brassica rapa* (Weis *et al.* 2015). In addition to highlighting an unexpected sensitivity of softness to genes, individuals and population densities, it was found in both cases that the softness of selection was generally high, cementing the idea that soft selection shapes natural variation at local adaptation loci (Agrawal 2010, Reznick 2016).

The present experimental work provides new perspectives for further testing theoretical predictions about the effect of spatial heterogeneity on polymorphism maintenance. The genetics of local adaptation used (two resistance alleles at two different loci) hampers the study of the emergence of local adaptation polymorphisms by gradual evolution. But the experimental system is likely relevant for questions regarding the maintenance of already existing polymorphisms, as, *e.g.*, species diversity in communities established in heterogeneous environments. For instance, one could test more systematically for the effect of habitat productivities and trade-off intensities on polymorphism maintenance. The present experiment conservatively considered full migration between the two habitats. But an important body of knowledge has explored the effect of migration between habitats on the conditions for polymorphism maintenance. Migration intensity (*e.g.,* Maynard Smith 1966), timing (*e.g.,* Ravigné *et al.* 2004, Débarre and Gandon 2011, Massol 2013), and bias (density dependent migration or habitat selection, *e.g.,* de Meeûs *et al.* 1993) affect the range of conditions favorable to polymorphism maintenance and could be tested through a similar experimental design. Estimating the relative importance of spatial and temporal variability of the environment in shaping polymorphism could also help our understanding of ecological specialization (Massol 2013).

## Acknowledgements

We are grateful to R. E. Lenski for providing *E. coli* strains. We thank C. Duperray (IRMB – Montpellier) and the Montpellier RIO Imaging platform for hosting the flow cytometry measurements, F. Débarre and B. Facon for helpful comments on the manuscript, and C. Prator for careful English editing. We thank three PCI Evolutionary Biology recommenders for their constructive corrections. This study received financial support by the French Agropolis Fondation (Labex Agro-Montpellier, BIOFIS Project Number 1001-001 and E-SPACE project number 1504-004), the European Union (ERDF), and the ‘Conseil Régional de La Réunion’.

## Authorship Statement

RF had the original idea of the study; RF, RG and VR designed the experiments; RF and RG carried out the experiments; RG analyzed experimental results; VR provided theoretical predictions; RF, RG and VR wrote the paper.

## Data Accessibility Statement

The data supporting our results will be archived in an appropriate public repository (Dryad/Agritrop) and the data DOI will be included at the end of the article, upon acceptance on the present manuscript.

## Competing Financial Interests Statement

The authors declare no competing financial interests.

## Supplementary Materials

**Table S1.**
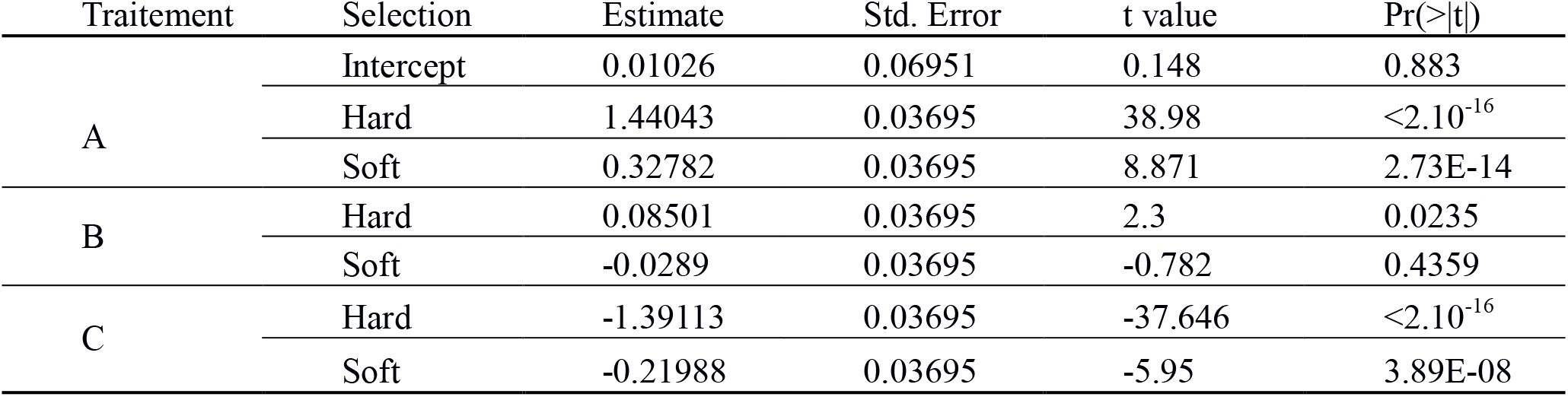
Statistical analysis of the relative genotype frequency variations. Relative frequencies were transformed with a logit function, in order to linearize the curves and therefore perform linear regressions. R = 0.975.

| Traitement | Selection | Estimate | Std. Error | t value | Pr(>\|t\|) |
| --- | --- | --- | --- | --- | --- |
| A | Intercept | 0.01026 | 0.06951 | 0.148 | 0.883 |
| | Hard | 1.44043 | 0.03695 | 38.98 | $<2.10^{-16}$ |
|  | Soft | 0.32782 | 0.03695 | 8.871 | 2.73E-14 |
| B | Hard | 0.08501 | 0.03695 | 2.3 | 0.0235 |
|  | Soft | -0.0289 | 0.03695 | -0.782 | 0.4359 |
| C | Hard | -1.39113 | 0.03695 | -37.646 | $<2.10^{-16}$ |
|  | Soft | -0.21988 | 0.03695 | -5.95 | 3.89E-08 |

**Figure S1.**
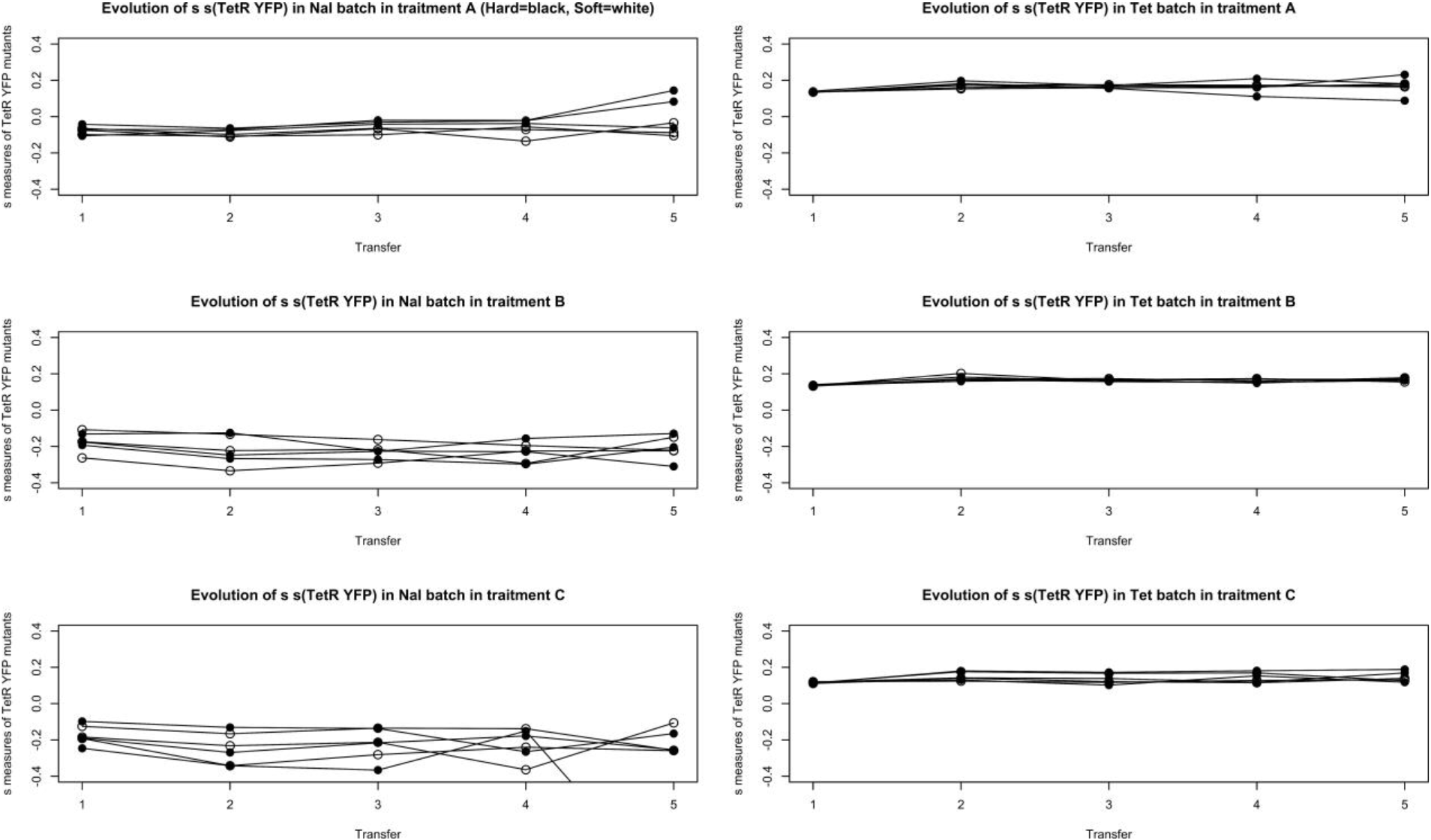
Evolution of selection coefficients (*s*) during the experiment. Graphs on the left side show the evolution of *s* in the Nal-habitat, while graphs on the right side show the evolution of *s* in the Tet-habitat. It is to be noted that when relative frequencies reach extreme values (close to 0 or 1), the estimation of *s* are less precise, due to the detection of only few individuals of the losing genotype. This explains some s variations at transfer 5 in treatments A and C, and more specifically why we observed a very low *s* measure in the Nal habitat at transfer 5, in one population in treatment C.

## References

1. Agrawal, A.F. (2010). Ecological Determinants of Mutation Load and Inbreeding Depression in Subdivided Populations. Am. Nat., 176, 111–122

2. Bell, G. (1997). Experimental evolution in *Chlamydomonas.* I. Short-term selection in uniform and diverse environments. Hered., 78, 490–497

3. Bell, G. and Reboud, X. (1997). Experimental evolution in *Chlamydomonas.* II. Genetic variation in strongly contrasted environments. Hered., 78, 1–11

4. Chao, L., Hanley, K. a, Burch, C.L., Dahlberg, C. and Turner, P.E. (2000). Kin selection and parasite evolution: higher and lower virulence with hard and soft selection. Q. Rev. Biol., 75, 261–275

5. Chevin, L.-M. (2011). On measuring selection in experimental evolution. Biol. Lett., 7, 210–3

6. Christiansen, F.B. (1974). Sufficient conditions for protected polymorphism in a subdivided population. Am. Nat., 108, 157–166

7. Christiansen, F.B. (1975). Hard and Soft Selection in a Subdivided Population. Am. Nat., 109, 11–16

8. Crandall, K. a, Bininda-emonds, O.R.P., Mace, G.M. and Wayne, R.K. (2000). Considering evolutionary pressures in conservation biology. Trends Ecol. Evol., 15, 290–295

9. Datsenko, K. a and Wanner, B.L. (2000). One-step inactivation of chromosomal genes in *Escherichia coli* K-12 using PCR products. Proc. Natl. Acad. Sci. U. S. A., 97, 6640–6645

10. De Meeûs, T., Michalakis, Y., Renaud, F. and Olivieri, I. (1993). Polymorphism in heterogeneous environments, evolution of habitat selection and sympatrci speciation - soft and hard selection models. Evol. Ecol., 7, 175–198

11. Débarre, F. and Gandon, S. (2011). Evolution in Heterogeneous Environments : Between Soft and Hard Selection. Am. Nat., 177, E84–E97

12. Dempster, E.R. (1955). Maintenance of genetic heterogeneity. Cold Spring Harb. Symp. Quant. Biol. Sci., 20, 25–32

13. Ebert, D. (1998). Experimental evolution of parasites. Science, 282, 1432–1436

14. Gallet, R., Cooper, T.F., Elena, S.F. and Lenormand, T. (2012). Measuring selection coefficients below 10^-3^: Method, Questions, and Prospects. Genetics, 190, 175–186

15. García-Dorado, A., Martin, P. and García, N. (1991). Soft selection and quantitative genetic variation: a laboratory experiment. Heredity, 66, 313–323

16. Gigord, L.D.B., Macnair, M.R. and Smithson, A. (2001). Negative frequency-dependent selection maintains a dramatic flower color polymorphism in the rewardless orchid *Dactylorhiza sambucina*(L.) Soo. Proc. Natl. Acad. Sci., 98, 6253–6255

17. Ho, E.K.H. and Agrawal, A.F. (2012). The effects of competition on the strength and softness of selection. J. Evol. Biol., 25, 2537–2546

18. Hori, M. (1993). Frequency-Dependent Natural Selection in the Handedness of Scale-Eating Cichlid Fish. Science, 260, 216–219

19. Jasmin, J.N. and Kassen, R. (2007). On the experimental evolution of specialization and diversity in heterogeneous environments. Ecol. Lett., 10, 272–281

20. Johnson, D.W., Christie, M.R., Moye, J. and Hixon, M.A. (2011). Genetic correlations between adults and larvae in a marine fish: Potential effects of fishery selection on population replenishment. Evol. Appl., 4, 621–633

21. Juenger, T., Lennartsson, T. and Tuomi, J. (2000). The evolution of tolerance to damage in *Gentianella campestris*: Natural selection and the quantitative genetics of tolerance. Evol. Ecol., 14, 393–419

22. Karlin, S. and Campbell, R.B. (1981). The existence of a protected polymorphism under conditions of soft as opposed to hard selection in a multideme population system. Am. Nat., 117, 262–275

23. Kassen, R. (2002). The experimental evolution of specialists, generalists, and the maintenance of diversity. J. Evol. Biol., 15, 173–190

24. Kelley, J.L., Stinchcombe, J.R. and Weinig, C. (2005). Soft and hard selection on plant defense traits. Evol. Ecol. Res., 287–302

25. Kleckner, N., Bender, J. and Gottesman, S. (1991). Uses of transposons with emphasis on Tn10. Methods Enzymol., 204, 139–180

26. Laffafian, A., King, J.D. and Agrawal, A.F. (2010). Variation in the strength and softness of selection on deleterious mutations. Evolution., 64, 3232–3241

27. Lenski, R.E., Rose, M.R., Simpson, S.C. and Tadler, S.C. (1991). Long-Term Experimental Evolution in *Escherichia coli.* I. Adaptation and Divergence During. Am. Nat., 138, 1315–1341

28. Levene, H. (1953). Genetic equilibrium when more than one niche is available. Am. Nat., 87, 331–333

29. Levins, R. (1962). Theory of fitness in heterogeneous environments. 1. The fitness set and the adaptive function. Am. Nat., 96, 361–373

30. Mackauer, M. (1990). Host discrimination and larval competition in solitary endoparasitoids. In: Critical Issues in Biological Control (eds. Mackauer, M., Ehler, L.E. and Roland, J.). Intercept, Andover, pp. 41–62

31. Massol, F. (2013). A framework to compare theoretical predictions on trait evolution in temporally varying environments under different life cycles. Ecol. Complex., 16, 9–19

32. Maynard Smith, J. (1966). Sympatric Speciation. Am. Nat., 100, 637–650

33. Maynard, Smith, J. and Hoekstra, R. (1980). Polymorphism in a varied environment: how robust are the models? Genet. Res., 35, 45–57

34. Olendorf, R., Rodd, F.H., Punzalan, D., Houde, A.E., Hurt, C., Reznick, D.N., et al. (2006). Frequency-dependent survival in natural guppy populations. Nature, 441, 633–636

35. Prout, T. (1968). Sufficient Conditions for Multiple Niche Polymorphism. Am. Nat., 102, 493–496

36. Rainey, P.B., Buckling, A., Kassen, R. and Travisano, M. (2000). The emergence and maintenance of diversity : insights from experimental bacterial populations, Trends Ecol. Evol. 5347, 243–247

37. Ravigné, V., Dieckmann, U. and Olivieri, I. (2009). Live where you thrive: joint evolution of habitat choice and local adaptation facilitates specialization and promotes diversity. Am. Nat., 174, E141-69

38. Ravigné, V., Olivieri, I. and Dieckmann, U. (2004). Implications of habitat choice for protected polymorphisms. Evol. Ecol. Res., 6, 125–145

39. Reznick, D. (2016). Hard and Soft Selection Revisited : How Evolution by Natural Selection Works in the Real World. J. Hered. 3–14

40. Sinervo, B. and Lively, C. (1996). The Rock-paper-scissors game and the evolution of alternative male strategies. Nature, 380, 240–243

41. Thrall, P.H., Oakeshott, J.G., Fitt, G., Southerton, S., Burdon, J.J., Sheppard, A., et al. (2011). Evolution in agriculture: the application of evolutionary approaches to the management of biotic interactions in agro-ecosystems. Evol. Appl., 4, 200–215

42. Vale, P.F. (2013). Killing them softly: Managing pathogen polymorphism and virulence in spatially variable environments. Trends Parasitol., 29, 417–422

43. Wade, M.J., Bijma, P., Ellen, E.D. and Muir, W. (2010). Group selection and social evolution in domesticated animals. Evol. Appl., 3, 453–465

44. Wallace, B. (1975). Hard and soft selection revisited. Evolution, 465-473

45. Weis, A.E., Turner, K.M., Petro, B., Austen, E.J. and Wadgymar, S.M. (2015). Hard and soft selection on phenology through seasonal shifts in the general and social environments: A study on plant emergence time. Evolution, 69, 1361–1374

46. Whitlock, M.C. (2002). Selection, Load and Inbreeding Depression in a Large Metapopulation, Genetics, 1202, 1191–1202

